# Repurposing The Dark Genome. III - Intronic Proteins

**DOI:** 10.1101/2023.06.10.544447

**Authors:** Mohit Garg, Pawan K. Dhar

## Abstract

Based on the expression patterns, genomes are viewed as a collection of protein-coding, RNA-coding, and non-expressing DNA sequences. Unlike most prokaryotes, eukaryotic gene expression comes with an additional step called alternative splicing. During the maturation process, different combinations of exons are spliced out and joined together resulting in the formation of mRNA isoforms. After removal from pre-mRNA, introns may be degraded by cellular exonucleases or form long non-coding RNAs (lncRNAs), or temporarily retained in the nucleus for regulating gene expression. We asked: Do introns have an unutilized potential for encoding proteins? If introns had an opportunity of getting translated, what kind of peptides or proteins, would they make? This study is based on the hypothesis of making functional proteins from leftover introns and is an extension of the original work of making functional proteins from the *E. coli* intergenic sequences (Dhar et al., 2009). Here full-length introns were computationally translated into proteins to study their potential structural, physicochemical, functional, and cellular location properties. Experimental validation is underway for a detailed understanding of the biology of intronic proteins. A synthetic intronic protein repository would provide an opportunity to design first-in-the-class molecules toward functional endpoints.

## 1. Introduction

A genome is the complete set of genetic material, including all the genes and non-expressing DNA sequences, present in an organism. The genome provides the instructions for building and maintaining the organism, determining its characteristics and traits. Based on their functional patterns, genomes are viewed as a collection of protein-coding, RNA-coding, and non-expressing sequences.

Unlike most prokaryotes, in eukaryotes, transcription occurs in two steps. The first step consists of splicing out of introns from pre-mRNA and the second step is joining exons of pre-mRNA to form a mature mRNA ready for translation.

For a long time, it was believed that post-splicing, introns were sent for degradation and possibly recycled. However, several compelling pieces of evidence generated a new thought of introns getting reused for regulatory roles post-splicing (Hesselberth, 2013).

Currently, the view is that the fate of introns is organism-specific. Some of the common intron disposal channels are (i) degradation by exonucleases to ensure that introns do not interfere with the translation of mRNA (ii) recycling of intronic sequences in the form of long non-coding RNAs (lncRNAs), microRNAs, and so on (Jo & Choi, 2015) and (iii) temporary retention of introns in the nucleus to fulfill a certain regulatory role.

However, to our best knowledge, currently, there is no report of introns getting translated into polypeptides in their natural setting.

Here, we have addressed this unutilized potential of introns and attempted to generate a possibility of making synthetic peptides or proteins from intronic sequences that are naturally programmed to make cellular debris.

The concept of making functional protein molecules from non-expressing DNA (dark genome) came in 2009 when experimental evidence of synthetically making growth-inhibitory proteins from intergenic sequences of *E. coli* was provided (Dhar et al., 2009).

## 2. Material and Methods

The *Saccharomyces cerevisiae* (yeast) intron database and *Caenorhabditis elegans* intron database, and *Drosophila melanogaster* (fruit fly) intron database were used in this study.

The data was downloaded from The Intron Annotation and Orthology Database (IAOD) (Moyer et al., 2019). A total of 332 Introns from *S. cerevisiae*, 1,10,321 introns from *C. elegans*, and 46,800 introns from *D. melanogaster* genomes were used for the study shown in Fig. 1. The overall methodological approach was the same as reported earlier (Garg & Dhar, 2023).

**Fig. 1:**
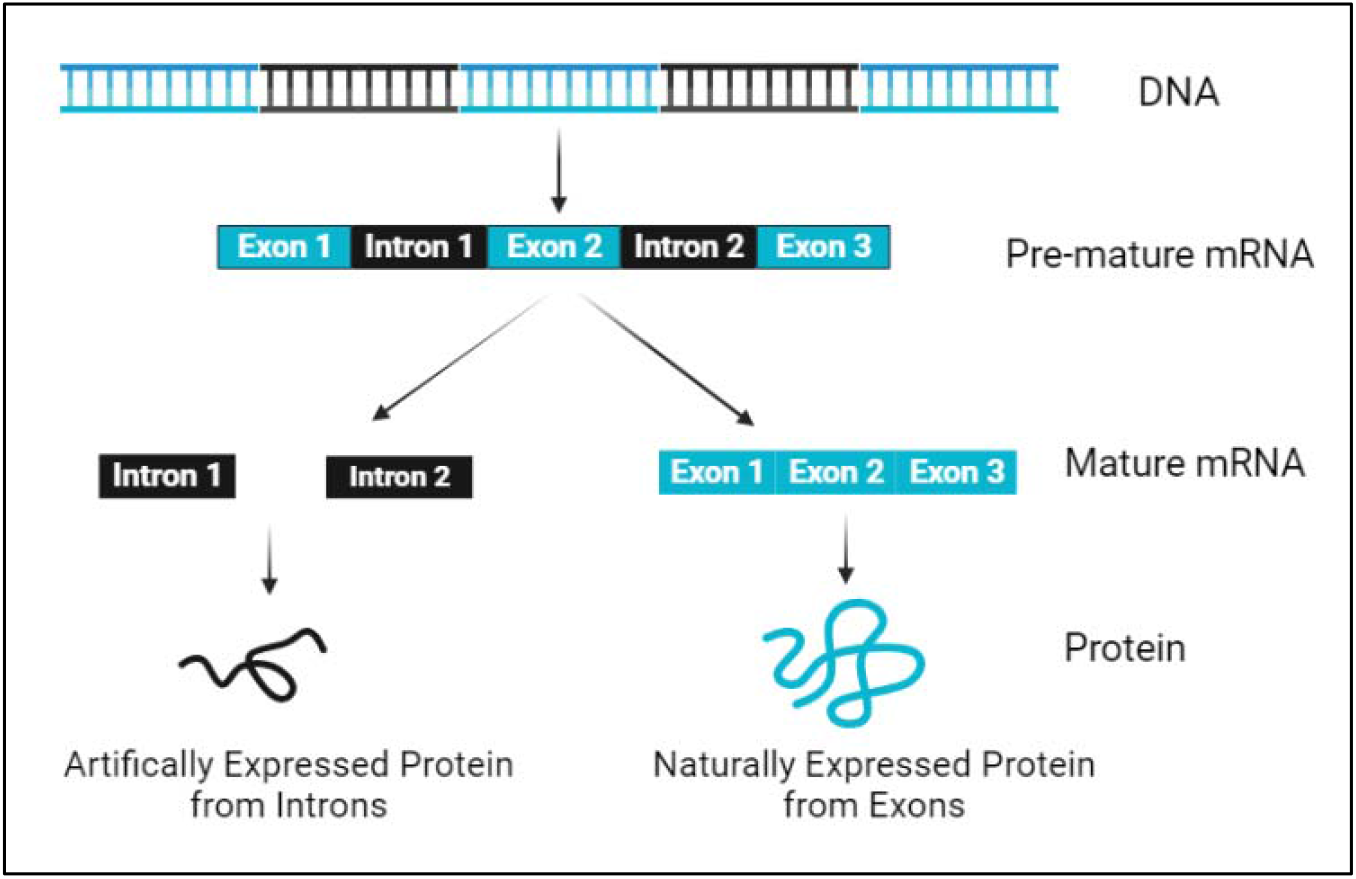
Schematic diagram for making proteins from the genes. Both naturally expressing proteins from exons and artificially expressing proteins from introns.

### 2.1 Translation of Intron sequences

DNA Sequences of introns of the *S. cerevisiae, C. elegance*, and *D. melanogaster* were computationally translated into protein/peptide sequences using the UGENE tool (Okonechnikov et al., 2012). Only the full-length translates were used i.e., those that did not show any stop codon on translation. The quality check of the translation is done by the Expasy Translation tool (Gasteiger, 2003).

### 2.2 Protein Sequence Similarity

Full-length translated sequences were BLASTed against a previously available protein database (Non-Redundant Protein Database) in the NCBI Blast Server (Altschul et al., 1990). Sequences that did not show any similarity with the existing protein sequence database were included for further study.

### 2.3 Physicochemical properties

The Expasy-ProtParam tool of SIB Swiss Institute of Bioinformatics (Roy et al., 2011) was used to analyze physiochemical properties of sequences such as molecular weight, theoretical pI, instability index, and grand average of hydropathicity (GRAVY) index. The physicochemical properties indicate the characteristics and nature of proteins.

### 2.4 Structure and Function Prediction

The structure and Function of protein sequences were predicted by the I-TASSER Server (Iterative Threading ASSEmbly Refinement) tool (Zhang, 2008). Secondary and tertiary structures are predicted by I-TASSER, along with probable functions. The WoLF PSORT tool (Horton et al., 2007)was used to analyze the protein’s subcellular localization.

### 2.5 Stereochemical properties of protein models

The protein model’s stereochemical properties were analyzed using Procheck Server (Laskowski et al., 1993). The Ramachandran plot assessed the geometry of amino acids. (Hollingsworth & Karplus, 2010).

## 3. Results

### 3.1 Translation of Intron sequences

Computational translation of intron DNA sequences generated many full-length sequences (Table-1).

**Table 1:**
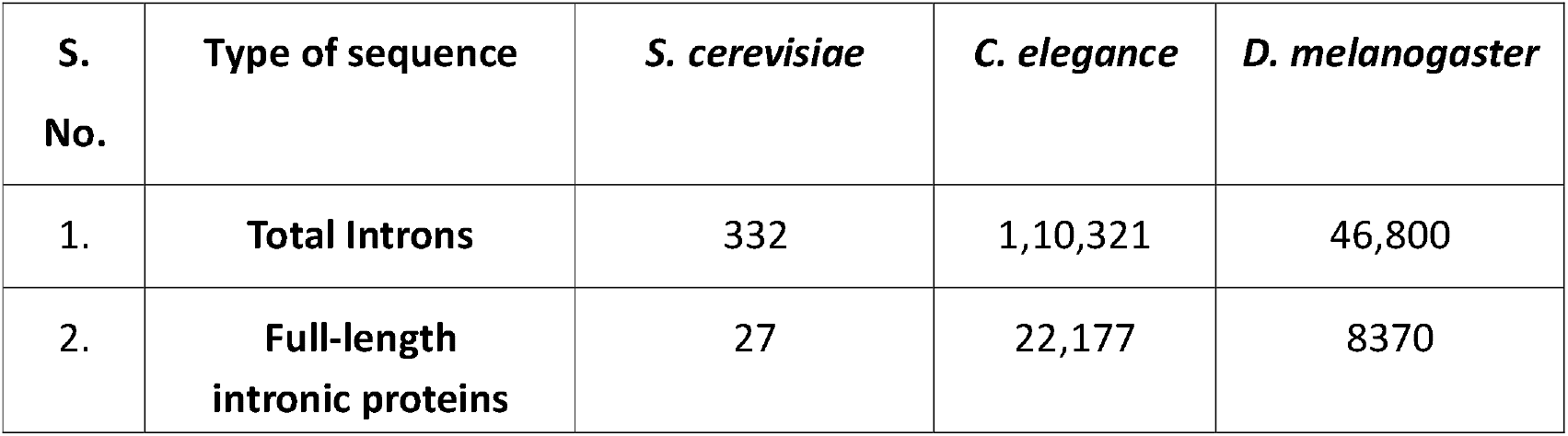
Full-length translated introns sequences in *S. cerevisiae, C. elegance*, and *D. melanogaster*

| S. No. | Type of sequence | *S. cerevisiae* | *C. elegance* | *D. melanogaster* |
| --- | --- | --- | --- | --- |
| 1. | Total Introns | 332 | 1,10,321 | 46,800 |
| 2. | Full-length intronic proteins | 27 | 22,177 | 8370 |

### 3.2 Protein Sequence Similarity

Full-length Intron sequences of *S. cerevisiae, C. elegance*, and *D. melanogaster* were BLASTed against the NCBI protein sequences database. A subset of sequences did not show any sequence similarity with the known proteins.

In this study, we considered a random subset of full-length translates from *S. cerevisiae, D. melanogaster*, and *C. elegance* genomes.

### 3.3 Physicochemical properties

The Intron-encoded proteins were studied for physiochemical properties (Table 2). The molecular Weight of intronic proteins was found to be in the range from 3.8 kDa to 17.8 kDa, with an Isoelectric Point (pI) value ranging from 4.30 to 11.11 (pI<7 indicating the basic nature of intronic proteins), Instability index ranging from 5.35 to 44.80 (indicating more stable structure), an aliphatic index ranging from 57.80 to 150.30 (indicating thermal stability), hydropathicity value (GRAVY) in the range of -0.46 to 2.05 showing interaction with water (low GRAVY score indicates better interaction with water), and G-C ratio ranging from 28.6 – 57.1.

**Table 2.** Physiochemical Properties of introns proteins of *S. cerevisiae, C. elegance*, and *D. melanogaster*

| S. No. | Physiochemical properties | *S. Cerevisiae* | *C. Elegance* | *D. Melanogaster* |
| --- | --- | --- | --- | --- |
| 1. | Length - Amino Acid (range) | 32 - 65 | 86 - 108 | 80 - 164 |
| 2. | Molecular weight (range) (kDa) | 3.8 - 7.8 | 9.8 - 12.2 | 9.1 - 17.8 |
| 3. | Theoretical pI (range) | 4.30 - 10.03 | 5.43 - 11.11 | 7.74 - 10.5 |
| 4. | Instability index (range) | 14.60 - 40.70 | 5.35 - 24.64 | 22.90 - 44.80 |
| 5. | Aliphatic index (range) | 57.80 - 150.30 | 70.50 - 113.8 | 61.27 - 116.79 |
| 6. | Grand average of hydropathicity (GRAVY) (range) | -0.46 to 2.052 | -0.51 to 0.593 | -0.607 to 0.618 |
| 7. | G-C Ratio (range) | 28.6 - 37.7 | 30.5 - 57.1 | 40.4 - 56.1 |

### 3.4 Structure and Function Prediction

The structure and function of intronic proteins predicted by I-Tasser showed a favorable C-score, i.e., within the range of -5to+2. The structural similarity of proteins ranges from 0.456 to 0.881(Table 3). In the predicted structure of proteins, we observed that introns protein shows natural folding (α-Helix and β-sheets) like naturally expressing proteins (Fig. 2). Subcellular localization of proteins predicted by WoLFPSORT showed primary localization of proteins into mitochondria (inner membrane and matrix), different places in cytoplasm included with the nucleus, and some proteins found as secretory proteins. Predicted Intronic proteins show binding with other materials such as organic compounds, Chlorophylla, and Ions like calcium, Zinc, and Manganese (Table 4).

**Table 3.**
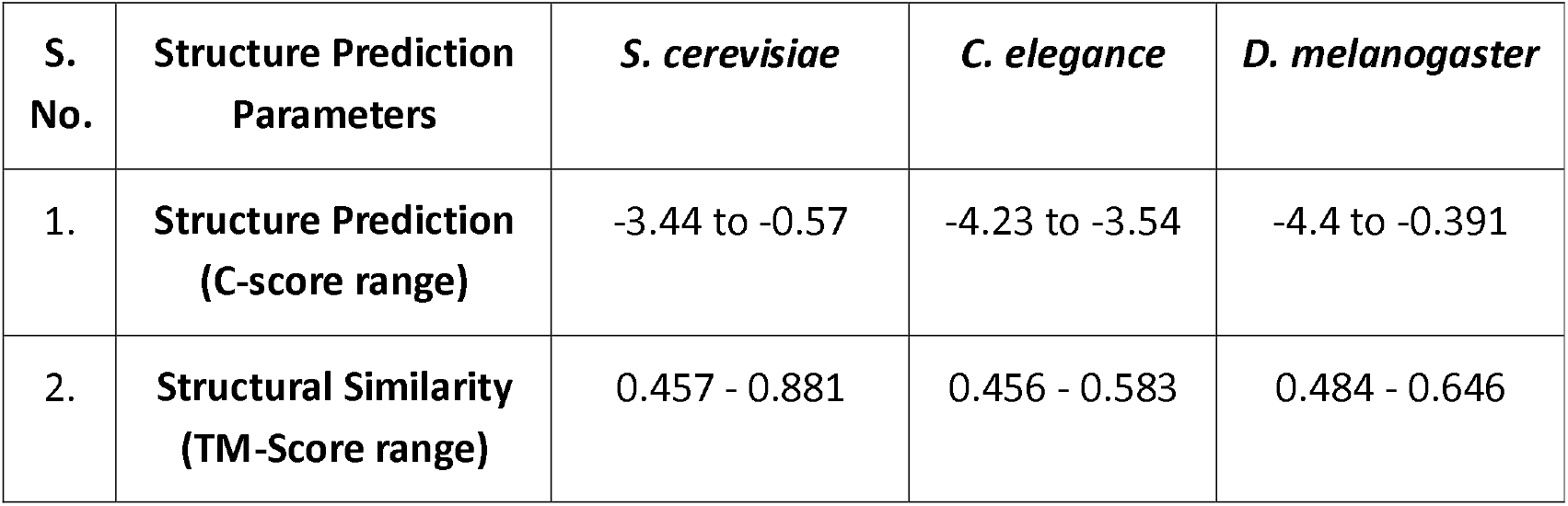
Predicted Structure and structural similarity of introns proteins of *S. cerevisiae, C. elegance*, and *D. melanogaster*

| S. No. | Structure Prediction Parameters | *S. cerevisiae* | *C. elegance* | *D. melanogaster* |
| --- | --- | --- | --- | --- |
| 1. | Structure Prediction (C-score range) | -3.44 to -0.57 | -4.23 to -3.54 | -4.4 to -0.391 |
| 2. | Structural Similarity (TM-Score range) | 0.457 - 0.881 | 0.456 - 0.583 | 0.484 - 0.646 |

**Table 4.** Function and Subcellular Localisation prediction of introns proteins of *S. cerevisiae, C. elegance*, and *D. melanogaster*

| S. No. | Predictions | *S. cerevisiae* | *C. elegance* | *D. melanogaster* |
| --- | --- | --- | --- | --- |
| 1. | Function prediction (types) | ligase, transferase, transcription Inhibitor, transport protein, oxidoreductase | ligase, transferase, lyase, DNA binding protein, Cell adhesion protein, oxidoreductase | photosynthesis, membrane protein, signaling protein, hydrolase, transferase, isomerase |
| 2. | Ligand binding prediction (Types) | organic compounds, calcium, zinc Ions | organic compounds, zinc copper ions | organic compounds, manganese ions |
| 3. | Subcellular Localisation | cytoplasmic, Integral membrane protein, secretory | mitochondrial, nuclear, cytoplasmic | cytoplasmic, mitochondria, secretory |

**Fig. 2.**
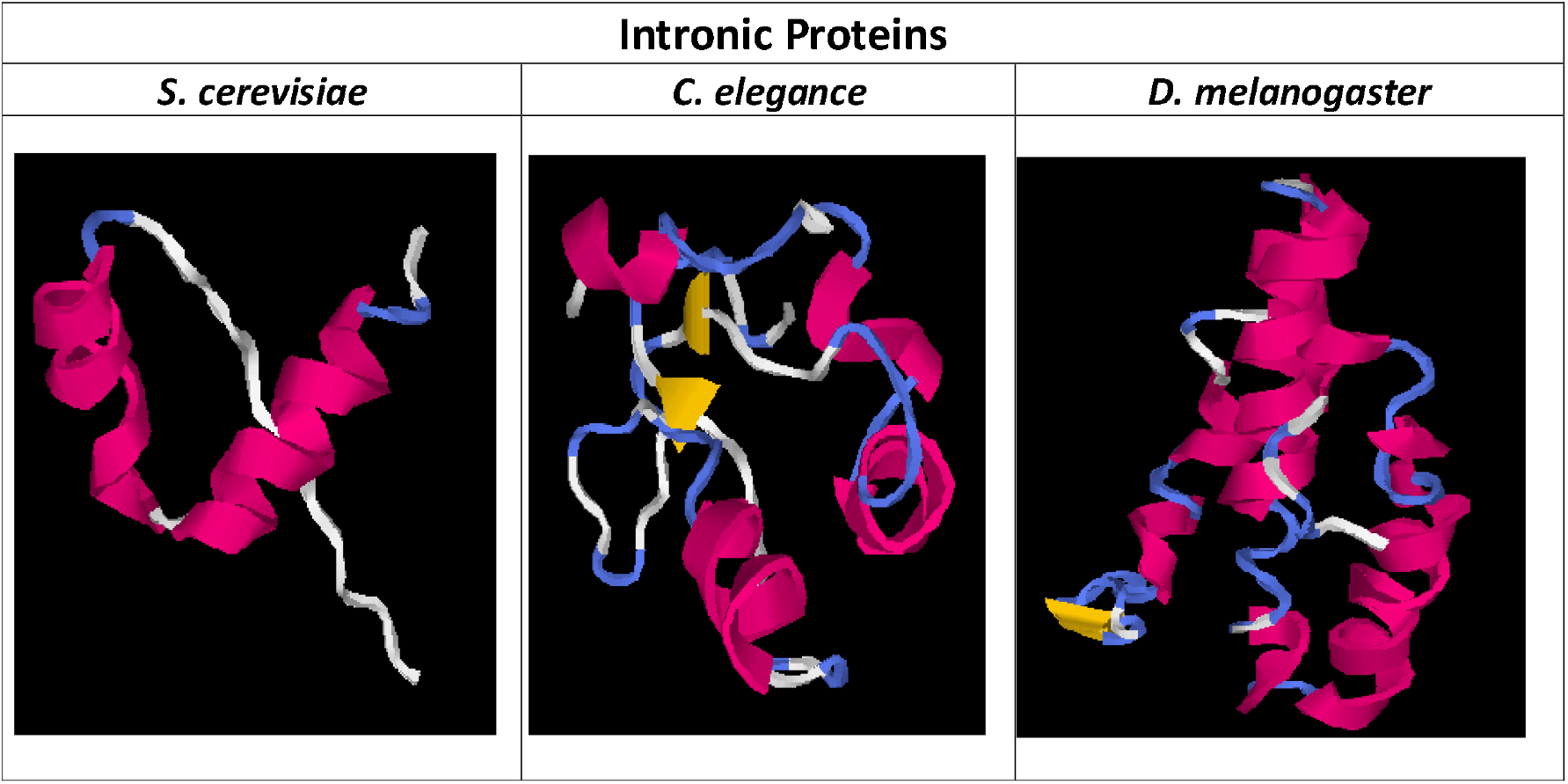
The Predicted tertiary structure of intronic protein sequences from *S. cerevisiae, C. elegance* and *D. melanogaster* resembling naturally expressing proteins

### 3.5 Stereochemical properties of protein models

Ramachandran plots of proteins were derived from studying the stereochemical properties of proteins (Fig. 3). Most of the amino acid residue of proteins appeared in the favorable regions (α-Helix and β-sheets) in the plot, indicating a promising folding profile of proteins.

**Fig. 3.**
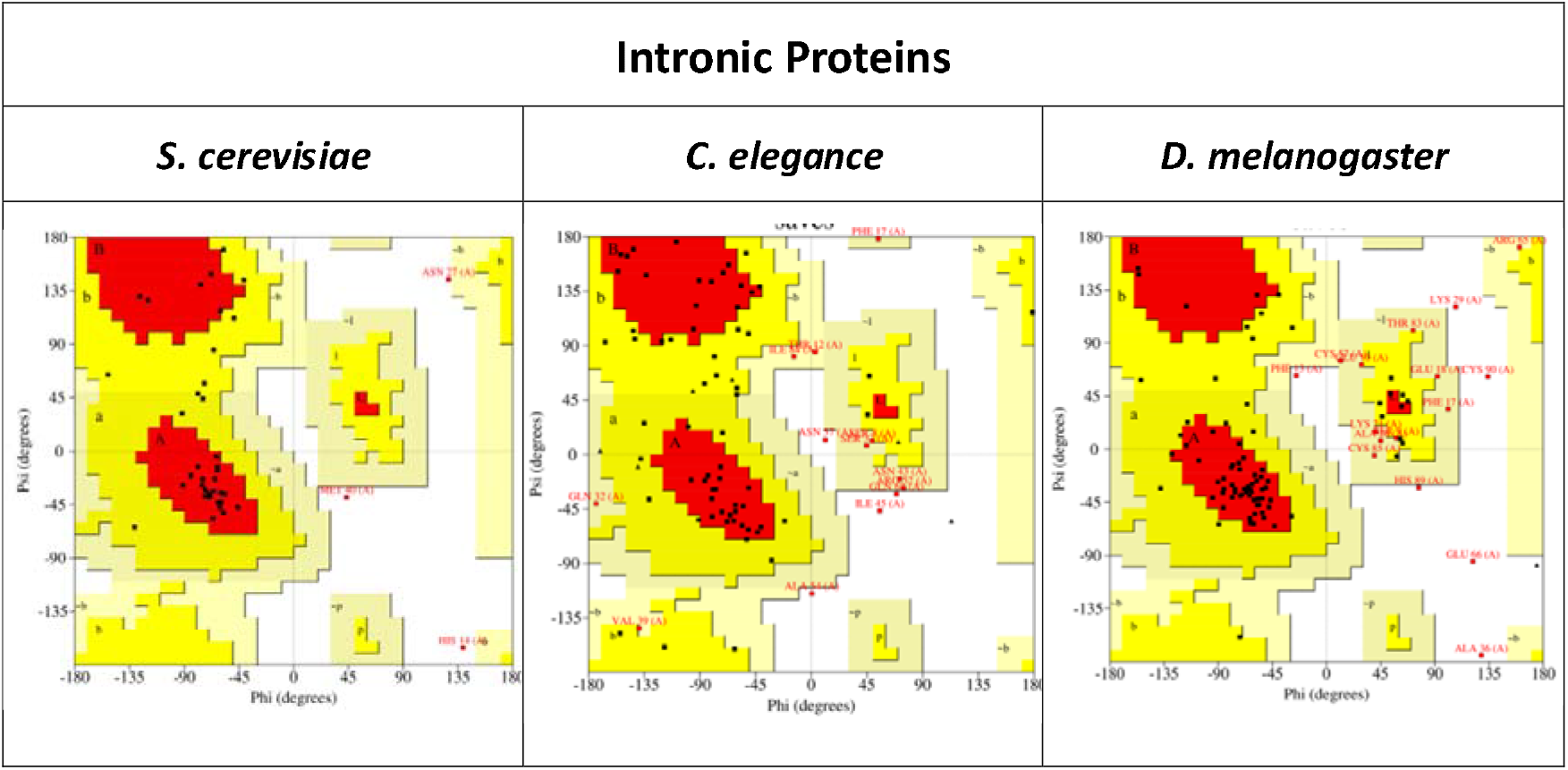
Ramachandran plot of intronic protein sequences of *S. cerevisiae, C. elegance*, and *D. melanogaster*. Most amino acid residues were found in favorable regions (red and dark yellow), indicating good protein folding.

## 4. Discussion

Introns are segments of DNA interspersed between coding regions, known as exons. Intronic sequences are primarily found in eukaryotic organisms, including plants, animals, and fungi.

Before translation, introns are removed in a process called RNA splicing. The remaining exons are joined together to form mature mRNA molecules for subsequent translation at the ribosomal interface. The presence of introns adds a layer of proteomic variation and regulatory complexity. Some of the potential functions of introns include regulating gene expression, providing evolutionary advantages, facilitating DNA recombination, and protecting coding sequences from mutations.

Though reports are hinting at the possibility of intronic proteins in some cells, a deliberate process of synthetically designed functional peptides and proteins from introns seems to be novel.

Making functional peptides and proteins from introns of *S. cerevisiae, C. elegance*, and *D. melanogaster* is an extension of previous work where cell growth-inhibiting proteins were artificially made from intergenic sequences of *E. coli* (Dhar et al., 2009). Subsequently, computational studies indicated the possibility of making functional proteins from pseudogene (Shidhi et al., 2015).

Similarly, tRNA sequences of *E. coli* were expressed into tREPs showing strong anti-leishmania properties (Chakrabarti et al., 2022). Recently it was observed that antisense Sequences (Garg & Dhar, 2023) and reverse Sequences (Nayak & Dhar, 2023) can be artificially encoded into stable and functional protein molecules.

Here, we used computationally translated intron sequences of *S. cerevisiae, C. elegance*, and *D. melanogaster* to design full-length proteins. The intron-derived proteins showed a likelihood of folding into stable proteins.

The Ramachandran Plot showed most of the amino acid residue of intronic proteins falling in the most favorable regions. Intronic proteins are mapped to cellular addresses such as nuclear, mitochondria, and cytoplasm, with some being secretory proteins. Functional prediction of intronic-proteins showed up categories like transporter, membrane, and DNA-binding and enzymatic activity. Predictions indicated a possibility of calcium, zinc, and manganese ions as ligands for predicted proteins.

It would be interesting to experimentally synthesize intronic proteins and explore various functional outcomes.

## Author Contributions

PKD conceived the idea of making functional proteins from intronic DNA sequences, supervised the work, and wrote the final version of the manuscript. MG performed all the computational work and drafted the first version of the manuscript.

## Acknowledgment

MG would like to express its warmest thanks to Mr. Shubham Garg for helping with the key computational work and CSIR for providing NET-JRF Scholarship. PKD sincerely thanks JNU for the critical lab support and Prof. Binay Panda (JNU) for extremely helpful discussions.

